# Keratinizing Squamous and Intestinal Metaplasia in Urothelium of Long-term Catheterized Patients with Chronic Urinary Tract Infections

**DOI:** 10.64898/2026.09.09.750385

**Authors:** Harinder Singh, Ayan Roy, Syed M. Lokman, Nazifa Humaira, Rodrigo V. Eguez, Yanbao Yu, Rembert Pieper

## Abstract

**Purpose:** Many spinal cord-injured (SCI) patients are recurrently catheterized to manage urinary retention and incontinence. Microbial pathogens colonize urethral catheter surfaces, persist in form of biofilms and trigger inflammatory responses in the bladder. Our objective was to determine bladder tissue changes due to long-term inflammation using LC-MS proteomics.

**Experimental Design:** We examined longitudinal proteomic profiles from eight SCI patients, sampled over 2 to 6 months, in a patient-specific manner as well as compared with trauma patients diagnosed with acute urinary tract infections (UTI). The sample sources were urinary pellets (UP) from all subjects, and catheter biofilm extracts (CB) from SCI patients. Bioinformatic protein network and western blot analyses served to corroborate evidence of protein and cell differentiation changes in the urinary tract of SCI patients.

**Results:** More than 50 proteins with functional or structural roles in the epidermal cornified envelope were quantitatively increased in proteomic profiles from three SCI patients. This included stress response keratins and proteins involved in keratin crosslinking, e.g. transglutaminases and small proline-rich proteins, suggesting occurrence of keratinizing squamous epithelial metaplasia. Protein co-expression network analyses supported epithelial trans-differentiation changes. Nearly thirty proteins with gastro-intestinal epithelial cell expression specificity were enriched in proteomic profiles from a single SCI patient. Among those were mucin-2 and sucrase-isomaltase, secreted proteins detected in samples for only that patient via western blots. This finding was consistent with intestinal epithelial metaplasia.

**Conclusions and clinical relevance:** Without using invasive cystoscopy and histopathology, proteomic analyses from easily available UP and CB samples allowed us to infer cell differentiation changes in urothelial tissues of SCI patients with chronic infections, supporting a concept of clinical relevance for urothelial metaplasia screening, a precursor stage of urothelial neoplasms.

**Significance of Study:** This work has innovative value and biological significance in discovering abnormal mucosal epithelial trans-differentiation processes. Knowledge of metaplasia in catheterized SCI patients with chronically infected and inflamed urinary tracts is also clinically relevant. Metaplasia signatures were discovered from differential LC-MS proteomic analyses of urinary pellet and bladder catheter biofilm extracts obtained in longitudinally studies, without use of invasive medical procedures (cystoscopy). Distinct protein clusters were indicative of either keratinizing squamous epithelial metaplasia or intestinal metaplasia in urothelial tissues. Chronic inflammation is a known trigger of epithelial trans-differentiation, but intestinal metaplasia in urethral and bladder tissue has rarely been reported.

## Introduction

Recurrent long-term bladder catheterization is common medical practice to treat patients with neurogenic urinary bladder syndrome and other forms of urinary incontinence. [1,2] Among the patients who suffer from neurogenic bladders are those with spinal-cord injuries (SCI), multiple sclerosis, Parkinson’s disease, and stroke. [3] In the USA alone, more than 200,000 individuals live with SCI and require regular urological care. Key long-term complications of neurologic bladders are frequent infections of the lower urinary tract with pathogenic bacteria, especially in individuals using indwelling urethral catheters for bladder drainage. Rarer consequences are pyelonephritis, nephrolithiasis, urinary incontinence not responsive to therapy and bladder malignancies. [3] To avoid chronic infections, conservative medical care is catheter replacement every one or two weeks. Suprapubic cystostomy and surgical augmentation cystoplasty [2] are more invasive but alternative ways to moderate the risk of ascending renal infections and urosepsis. [4] The common microbial causes of catheter-associated urinary tract infections (CAUTI) are facultatively anaerobic bacteria that thrive in biofilms forming between external catheter surfaces and urethral or bladder mucosal walls: uropathogenic *Escherichia coli* (UPEC), *Klebsiella pneumoniae*, *Pseudomonas aeruginosa*, *Staphylococcus aureus*, *Enterococcus* and *Proteus spp*. [5–7] Urease allows some *Proteus spp.* to produce ammonia by metabolizing urea, which increases the urinary pH and causes precipitation of calcium and magnesium phosphate salt crystals in the lumen and on the surface of catheters. A biofilm matrix in which the pathogen, host cell debris and salts are embedded forms. [6] Several studies longitudinally examining indwelling urethral catheters placed in the urinary tract of patients for 1-3 weeks reported chronic colonization and infection with multiple bacterial pathogens. [8–10] While patients may not always have symptoms, immune cells infiltrate and tissue inflammation takes place. [10,11] CAUTIs, regularly treated with antibiotic drugs, aggravate the antimicrobial resistance problem, and inter-strain as well as inter-species transfers of antibiotic resistance genes reduce activity and modulate the targets of antibiotic drugs. [12,13]

Mass spectrometry-based proteomics enables surveys of pathogens and immune responses of the host to CAUTI as demonstrated in recent studies [10,14] and may lend itself to characterize long-term adaptations of mucosal tissues to CAUTIs. The terms trans-differentiation and metaplasia emerged to describe changes of one tissue into another. Two types of metaplasia identified in the urinary tract are keratinizing squamous cell metaplasia (KSM) and non-keratinizing squamous cell metaplasia. KSM features dedifferentiated umbrella cells and keratinized foci in urothelial tissues. [15–17] Clinical significance of these metaplasia types remains unclear. While considered reversible, metaplasia is also thought to be a precursor of urothelial hyperplasia and dysplasia that may progress to bladder malignancies. [18] Strong epidemiological links exist between chronic inflammation-associated schistosomiasis of the bladder, an infection caused by the protozoon *Schistosoma haematobium*, and squamous cell carcinoma (SCC) with long latencies. [19] Whether mechanisms leading to bladder cell carcinomas differ dependent on the pathogenic cause (viruses, *S. haematobium* or poly-microbial catheter biofilms) is unclear. [20] To gain more insights into urothelial tissues of long-term catheterized SCI patients we did not directly conduct histological analyses, but used a non-invasive route for patient urine and catheter specimen collections over up to six months and characterized their proteomes for inference of processes indicative of metaplasia.

## Methods

### 2.1. Ethics Statement

For human subject studies enabling collection of medical data and SCI patient specimens, a human subject protocol #56-RW-022 was approved by the Western Institutional Review Board in Olympia, Washington, and the J. Craig Venter Institute (JCVI) Review Board in Rockville, Maryland, in 2013. Use of patient specimens was designated ‘for research purposes only’. Upon signage of an informed consent form by visiting SCI patients, the Southwest Regional Wound Care Center in Lubbock, Texas, collected samples and medical data, assigning numerical codes to encrypt patient information prior to sharing those with lab scientists. Limited patient metadata were also published previously, with health risks assessed as minimal because sample collection was non-invasive, and catheter replacement was part of routine medical care. [10]

### 2.2. Study Design, Clinical Samples and their Preparation for LC-MS

Of nine enrolled adult participants, one patient dropped out of the clinical study fairly early. Longitudinally collected samples from eight SCI patients (6 male; 4 female) consisted of indwelling Foley catheters and urine specimens collected in catheter bags over 24 hours. [10] Catheter biofilm extract (CB) and urinary pellet (UP) samples, respectively, were donated over 2 to 6 months by the patients according to clinical appointment schedules. Matched UP and CB samples from a given timepoint were often, but not always, available. Shipments to the research site (JCVI) were arranged on dry ice overnight. Sample storage and cell extract preparation methods were described in [10]. Briefly, UP samples resulted from differential centrifugation (3,200 x g for 15 min) of pH-neutralized urine. CB samples were extracted in two steps: 1) agitating scraped biofilms with a solution of 100 mM sodium acetate (pH 5.5), 20 mM sodium meta-periodate and 300 mM NaCl, 2) recovering the soluble fraction centrifugally and 3) solubilizing cellular debris with a solution of 1% SDS (v/v), 5 mM EDTA and 50 mM DTT followed by 3-min heat treatment at 95°C. The two CB fractions were separately subjected to the LC-MS workflow. Protein contents were estimated from band staining intensities of ∼10 µl aliquots in CBB G250-stained gels. Aliquots of 100-200 µg protein were treated with denaturing solutions as described in the Filter-Aided Sample Preparation (FASP) protocol [21], with minor modifications: the total protein was digested with sequencing-grade trypsin at a 50:1 ratio (w/w) in Vivacon 10k filters (Sartorius AG, Germany) over 12-16 hours at 37°C. [22] Preparation of UP samples for LC-MS from the 18 trauma patients (with single timepoints) was similar and previously described. [11]

### 2.3. LC-MS/MS shotgun proteomic analysis

The LC-MS/MS workflow was described in [10]. Briefly, acidified aqueous acetonitrile LC gradients (195 min) were used to separate peptides from desalted FASP protein digests estimated to contain ∼5 µg peptides. The stationary phase was a PicoFrit C_18_ analytical column (75 μm × 15 cm, 3 μm pore size, 150 Å). The flow rate was set at 200 nl/min. The LC system was coupled via a FLEX nano-electrospray ion source to an LTQ-Velos Pro ion-trap mass spectrometer for acquisition of mass spectra applying 2.0 kV voltage distally, via a liquid junction. Peptide ions were analyzed in MS^1^ data-dependent mode, with the 10 most abundant positively charged fragment ions selected for MS^2^ scans. The MS duty cycles were controlled using XCalibur software v2.2, with collision-induced dissociation to fragment peptide ions set at a normalized collision energy of 35%. Dynamic exclusion was enabled. MS^1^ scans were selected from the m/z range of 380 to 1,800. Combined raw MS files from replicate LC-MS/MS runs were analyzed with the algorithm Sequest-HT, applying standard search parameters and FDR rates for rank 1 peptides that were estimated with the tool Percolator (versus a reverse sequence decoy database). Accepting protein hits equal to or lower than a 1% FDR threshold, protein groups were identified and quantified based on spectral-peptide match (PSM) counts. [22] The MS search database was composed of all non-redundant protein entries in the UniProt human proteome (release 2015-16) and 48 UniProt proteomes pertaining to microbial species selected for being a cause of UTIs. Only human protein IDs were relevant here. Raw and mzl files were deposited under data identifier PXD012048 in the public repository PRIDE (via ProteomeXchange). MS analysis of UP samples from the UTI-T cohort was performed on a Q-Exactive system (Thermo-Scientific). DDA and spectral data processing procedures and database searches were described in a previous publication. [11] LC-MS instruments and software tools were from Thermo-Scientific.

### 2.4. LC-MS/MS Data Normalization and Statistical Analysis

Protein IDs, the associated PSM counts, UniProt protein accession numbers, gene symbols, Mr values and unique peptide lists from all 126 MS datasets were integrated into a single data matrix. The PSM counts for each protein ID was normalized dividing by total human PSM counts in a given dataset and then log2-logerithmized. The protein log2 values are provided in data matrices included in Supplemental Material files. Protein lists were filtered by excluding those with missing values in ≥ 50% of all 126 datasets or in ≥ 33% of the dataset series from a single SCI patient. For principal component and hierarchical clustering analyses, 18 and 62 UP proteome datasets from the UTI-T and SCI cohorts, respectively, were used. Here, missing values were imputed with a function offered in Perseus v.2.1.3.0 software.[23] UP and CB datasets were also subjected to differential protein abundance analysis (DAA), with one exception separately, using unequal variance paired t-tests and correction for multiple testing via Benjamini-Hochberg calculations to determine statistical significance. Here, missing values were replaced by the lowest normalized PSM count observed. The groups for comparative analyses are defined in sections 3.4.-3.7. The significance threshold for P-values of the proteins was set at either ≤ 0.02 or ≤ 0.05 and the fold-change at ≥ 2. All Volcano plots were generated with a tool in the bioinformatics analysis software suite SRplot. [24]

### 2.5. Single Cell Type Expression, Protein Network and GO Term Enrichment Analyses

Using the Protein Atlas resource (https://www.proteinatlas.org/) we parsed records of unique or enriched tissue/single cell type expression. In the resource’s single cell tab, one section denotes ‘single cell type expression cluster’ entries. We used terms for such clusters as unbiased measures of a protein’s ‘specific cell type enrichment’. Several protein network analysis methods were applied. A simple one was the STRING bioinformatics resource tool (https://string-db.org/analysis) selecting the multi-protein tab. Refined multi-network analysis approaches [25,26] were employed to identify protein-specific co-expression modules comparing data from distinct patient groups. Protein co-expression network analysis (PCNA) [27] was performed using the Weighted Gene Co-expression Network Analysis tool, v1.72-1 in the R package. [28] Soft-thresholding power (β) to achieve an approximate scale-free topology (R^2^ ≥ 0.8) was 14. This was the lowest power to achieve a model fit with R^2^ greater than 0.8 (R^2^ = 0.88). Pairwise bi-weight mid-correlations (bicor) were used as an outlier-robust adjacency matrix, which was transformed into a topological overlap matrix for hierarchical clustering and dynamic tree cutting. Parameters were optimized for protein module detection as follows: deepSplit = 4, mergeCutHeight = 0.07, pamRespectsDendro = TRUE, reassignThreshold = p < 0.05, min ModuleSize = 12. Co-expressed protein modules were summarized by their first principal component (the module eigenprotein MEP) and correlated with a given SCI trait. Modules significantly associated with a trait (FDR < 0.05, r > 0.5) were prioritized. Key proteins were identified based on high module membership (kME > 0.8) and high protein significance (PS > 0.8 x module-trait correlation). [26] ClusterProfiler (v4.8.3) was used for functional enrichment analysis. [29] Over-representation of biological process GO terms was tested using Fisher’s exact test with Benjamini–Hochberg corrections (FDR < 0.05). The annotations were from database org.Hs.eg.db (v3.17.0). Bayesian inference analysis was applied to key proteins of selected modules using the BayesFactor package in R (v4.3.2), offering a superior alternative to the null hypothesis significance testing framework with evidence for proteome-phenotype associations. [30] A Bayes Factor (BF) > 10 was used as significance threshold. A 95% credible interval that does not overlap with zero [31] identified proteins strongly associated with a SCI patient trait.

### 2.6. Western blot experiments

Soluble fractions of UP and CB samples after heat-denaturation in a Tris-buffered 2.5% SDS and 2.5% β-mercaptoethanol solution were loaded onto SDS-PAGE mini-gels (4-12%T acrylamide) and run electrophoretically in MOPS buffer. One gel with nine samples was stained using CBB G-250 dye to adjust protein loading amounts at 5-15 µg per lane. Another gel with approximately equal total protein loads per lane was run followed by electroblotting onto a PVDF membrane sheet (at 100 Amps for 60-75 min). Transient staining with Ponceau S in 1% acetic acid was used to confirm satisfactory protein transfer. The blocking agent, primary antibody (Ab) dilutions (usually at 1:1000 in PBS-0.05% Tween-20), secondary antibody-enzyme conjugate dilutions (1:5000 in PBS-0.05% Tween-20), horseradish peroxidase (HRP) substrate concentration for chemiluminescent detection and time periods of PVDF membrane incubation steps were as reported previously. [32] Primary Abs were for mucin-2 (product no. sc-515032), sucrase-isomaltase #C-8 (product no. sc-393470), complement factor C3 (product no. EPR-19394) and cytokeratin-10 (product no. 111447). The latter Abs were from AbCam. The Abs with the prefix ‘sc’ and the antibody-enzyme conjugate (goat anti-rabbit IgG-HRP; product no. sc-2004) were obtained from Santa Cruz Biotechnology. HRP substrate exposure was stopped once luminescent protein bands began to emerge (maximally for 5 min). The ImageQuant LAS 500 Chemiluminescence CCD Camera was used for image capture.

## Results

### 3.1. Human subject cohorts and rationale for their comparative proteomic analysis

The main cohort consisted of SCI patients suffering from neuropathic bladder syndrome and required recurrent urethral catheter insertions to manage the urine voiding process. We examined catheterization time windows of 2 to 6 months for eight patients and collected between 6 to 9 UP and 5 to 8 CB samples from each patient. The biomasses of CB extracts typically represent a poly-microbial biofilm containing human cell secretions and debris, too. The longitudinal sampling design permitted the capture of patient-specific proteomic data repeatedly. All patients contracted CAUTIs, in nearly all cases the infections were persistent and resulted in chronic inflammation. [10] Here, our focus was on long-term perturbations of urothelial tissue not necessarily linked to antimicrobial defenses. For comparative purposes, UP proteomic profiles from 18 trauma patients diagnosed with acute UTIs (the UTI-T cohort) were included. The key differences (SCI vs. UTI-T) related to microbial colonization were: 1) *Proteus mirabilis* was a common, often dominant microbe in CAUTIs of SCI patients; 2) UPEC, known as the most prevalent cause of acute UTIs not associated with catheters, was indeed the most prevalent microbe in UTIs of UTI-T patients; 3) fastidious bacteria, which rarely cause UTIs/CAUTIs on their own, were identified as minor to moderately abundant microbes only in samples from SCI patients; 4) CAUTIs revealed higher bacterial diversity (richness).

### 3.2. Hierarchical clustering discerns SCI from UTI-T cohort and also separates SCI patient groups

Principal component analysis (PCA) allowed identification of patterns of the largest proteomic data variations along two orthogonal axes for the 80 UP datasets. As shown in ***Figure 1***, UTI-T data points largely separated from the longitudinally collected SCI data points. One SCI sample cluster featured many P1, P2, P7 and P8 data points (‘P’ for patient), another cluster was enriched for P4, P5, P6 and P9 data points (P5 had the most outliers). Then, we applied the Spearman and Pearson correlation distance matrices to hierarchically cluster all UP proteomic datasets. Spearman correlation analysis results along with a heat map are shown in ***Figure 2***, the Pearson correlation analysis is presented in Additional File 1 (Supplemental Materials). In ***Figure 2***, UTI-T sample identifiers begin with the term ‘d1’ or ‘d3’. SCI sample identifiers feature ‘P’ followed by the patient No. and a terminal single- or double-digit No. for a specific collection event. Hierarchical clustering revealed the separation of SCI from UTI-T cohorts. Furthermore, it depicts patient-specific sample clusters (6 to 8 collection events) and larger cluster branches (samples grouping two to four SCI patients). One group consisted of P1, P8, P2 and P7 datasets, another group of P6, P9, P4 and P5 datasets. Sub-groups we established were ‘P1-P8-P2’ and P7 that were chosen to perform DAAs compared with other datasets to gain insights into urothelial tissue adaptation processes (in sections ***3.4.***-***3.7.***). A complete set of UP proteomic data with protein lists, abundance values, normalization steps and statistical data assessments are presented in Additional File 2 (Supplemental Materials).

**Fig. 1.**
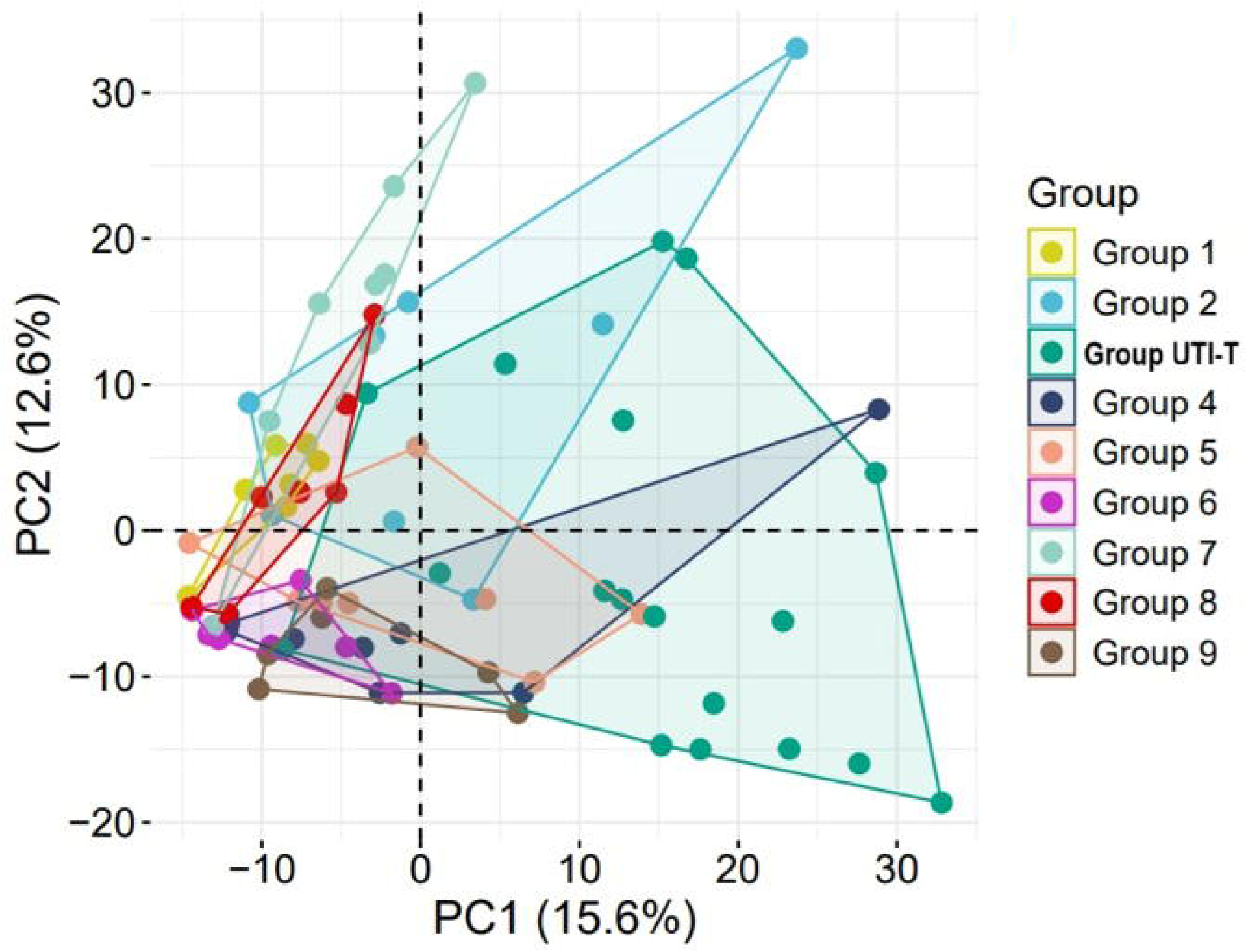
Principle component analysis of UP proteomic datasets derived from SCI and UTI-T cohorts. A matrix consisting of 18 UTI-T and 62 SCI datasets was uploaded into SRplot. The groups reflected longitudinal data from specific patients (P1 to P9) or data representing acute UTI (one sample/patient, UTI-T). Along the PC1 and PC2 axes, data represented the degree of variance the components captured from the original data.

**Fig. 2.**
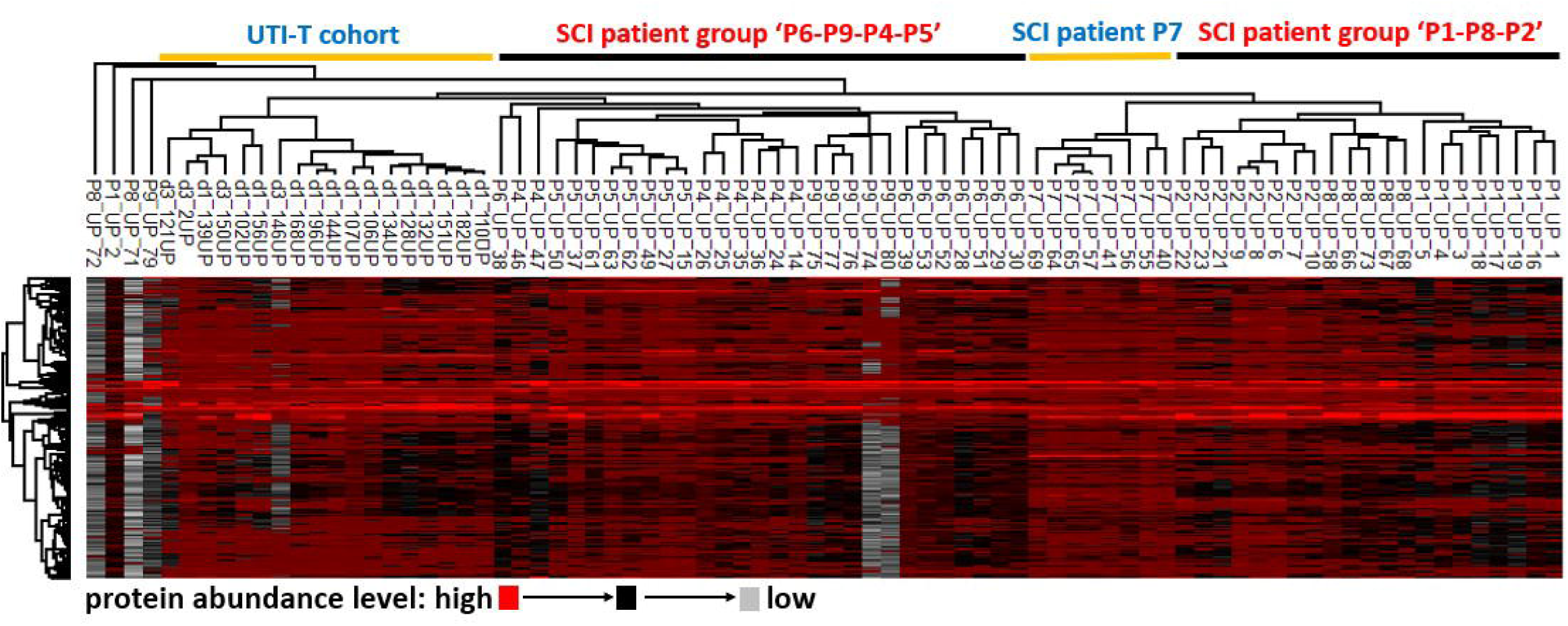
Hierarchical clustering using Spearman correlations to cluster samples and Pearson correlations to cluster proteins across 80 UP proteomic datasets. We note separate clustering of UTI-T samples and, largely, patient-specific SCI sample clustering. These results support the notion of underlying biological differences of the compared groups. The displayed dendrogram of samples explains our approach to select groups for differential abundance analyses (e.g., UTI-T, ‘P2-P8-P1’, ‘P6-P9-P4-P5’, P7). In Perseus v.2.1.3.0 software missing data were imputed. Only four major outliers among 80 samples in the clustering map were observed. With a distance threshold of 0.6, the analysis returned 36 protein clusters (dendrogram on the left). Protein abundances in the heat map are colored in a red-black-gray gradient (high to low).

### 3.3. Protein clustering for inferences of broad biological adaptations to CAUTI and acute UTI

Pearson correlation analysis was used to cluster proteins setting a distance threshold of 0.6, which returned 36 protein clusters (***Figure 2***). Zooming in, two clusters contained many proteins part of the acute phase response (APR). Dominant were components of the complement system. A third cluster featured many proteins highly expressed in granulocytes, especially neutrophils, and secreted into granules of named immune cells. These proteins and their assemblies are known for roles in the defense against UTI-causing pathogens. The 3 clusters are visualized in Additional Files 3 and 4a (Supplemental Materials). For cell type specific expression signatures, the Protein-Atlas data were resourced. Two proteins normally secreted by gastric and airway epithelial cells serving the local mucosal defense (BPIFB1, MUC5AC) formed another cluster, as shown in Additional File 4b. Two protein clusters we denote as keratin clusters KC1 and KC2 are shown in ***Figure 3***. Many proteins in KC1 and KC2 clusters are normally expressed at high levels only in keratinocytes of the epidermis and also squamous epithelia, but not in bladder epithelia. KC1 featured stress-responsive keratins (KRT-6a, KRT-6b, KRT-6c, KRT-17) and chaperonins active in keratinocytes (SFN and HSPB1). KC2 featured five small proline-rich proteins (SPRRs) that take part in the crosslinking of keratins and four proteins modulating both keratinocyte proliferation and fate (CALML3, S100A14, S100A16, CRNN). Each cluster depicted in ***Figure 4*** contained numerous proteins with enterocyte-specific expression patterns, therefore named intestinal clusters IC1 and IC2 here. While the heat map does not visualize this well due to missing value imputations, most proteins were exclusively detected in P7 datasets. IC1 featured the Fc-γ binding protein (FCGBP), chloride channel accessory protein 1 (CLCA1) and mucin-2 (MUC2), proteins with well-characterized roles in secretory goblet epithelial cells. IC2 contained 18 enterocyte-specific proteins. Examples are meprin-1 subunits A and B, cadherin CDHR5, intellectin-2 (ITLN2), REG3-α, the defensin DEFA5 and zymogen granule protein ZG16. Most of the proteins contribute to the maintenance of gut epithelial barrier integrity through structural roles or antimicrobial activities. Cluster IC2 also contained metabolic enzymes such as sucrase-isomaltase (SI) and nutrient transporters such as MTTP (for triglycerides). These proteins are specific to intestinal brush border cells (***Figure 4***).

**Fig. 3.**
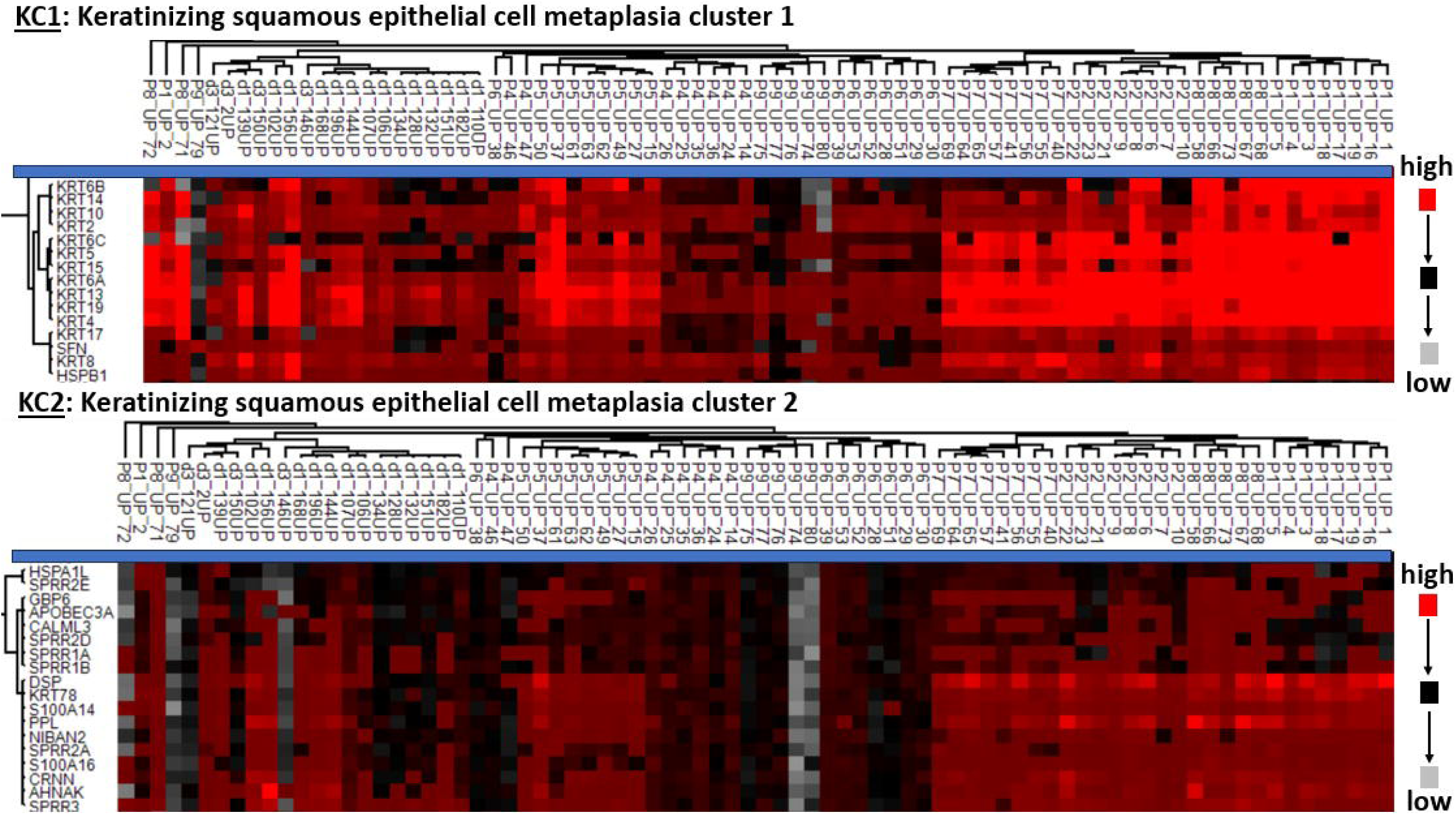
Protein clusters KC1 and KC2 (zoomed in on part of Fig.2 dendrogram/heat map) enriched in keratins and proteins involved in terminal keratinocyte differentiation and epidermal cornified envelope formation.

**Fig. 4.**
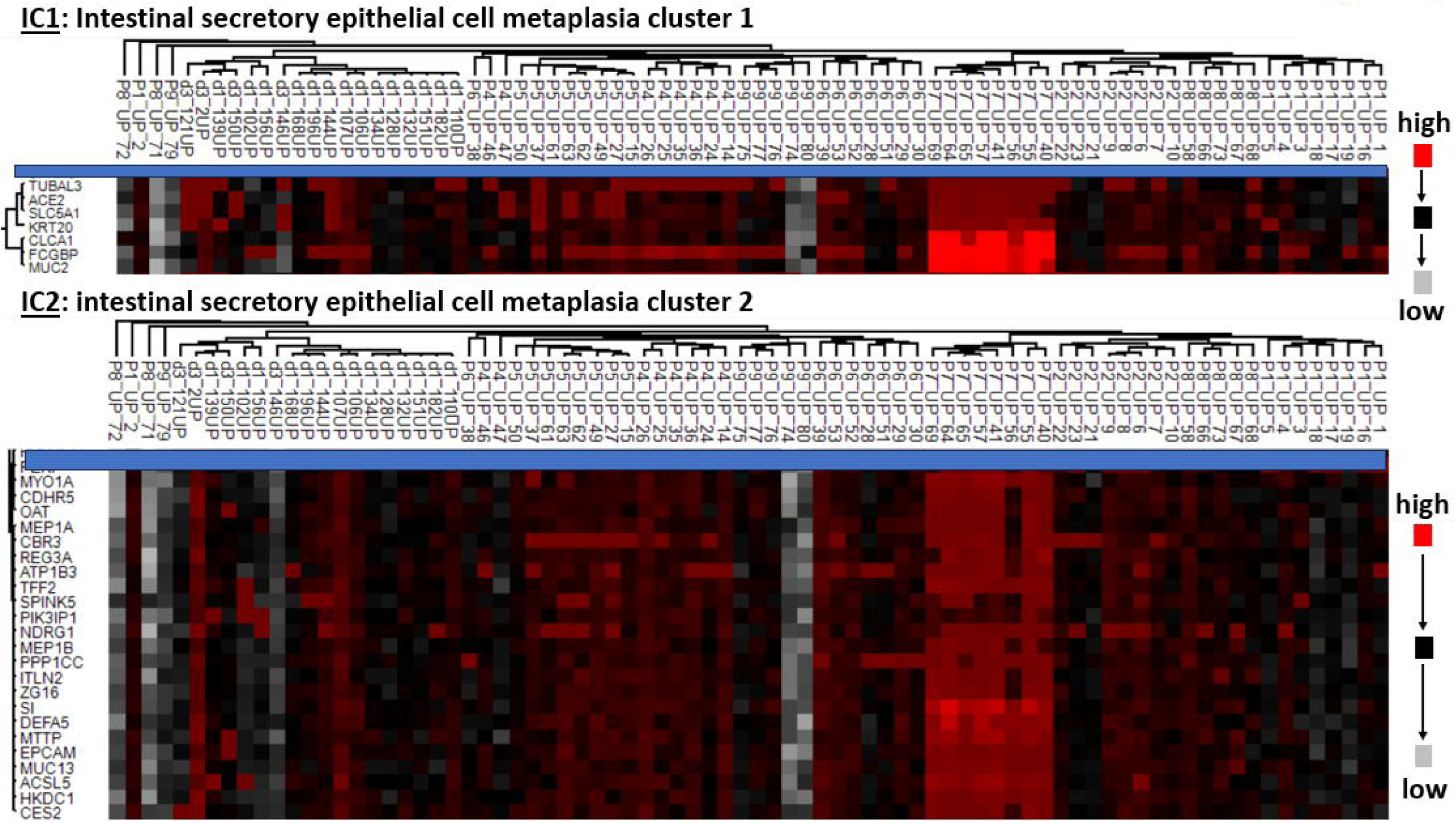
Protein clusters IC1 and IC2 (zoomed in on part of Fig.2 dendrogram/heat map) enriched in proteins normally unique to or enriched in expression in intestinal epithelial cells.

### 3.4. Keratinizing squamous cell metaplasia in three SCI patients is supported by DAA data

We identified the SCI patient group ‘P1-P8-P2’ from hierarchical clustering data (***Figure 1***). In unequal variance multiple testing-corrected t-tests (abbreviated ‘mtc•uevt-tests’ from here on), UP proteomic datasets of group ‘P1-P8-P2’ were compared to those of cohort UTI-T and those of SCI group ‘P6-P9-P4-P5’. Sixty proteins showed abundance increases in group ‘P1-P8-P2’ compared to cohort UTI-T, setting the threshold for statistical significance at P-value ≤ 0.02 and for fold-change at ≥ 2. Among those were 14 keratins including prominent type II/type I keratin pairs (KRT4/KRT13 and KRT5/KRT14), stress-responsive keratins (see *section 3.3*) as well as junction-plakoglobin (JUP) and desmoplakin (DSP), each a component of cell adherens junctions. All these proteins have roles in generating and modifying dynamic filament networks in stratified squamous epithelial cells. Also increased in abundance in group ‘P1-P8-P2’ vs. the UTI-T cohort were proteins involved in terminal keratinocyte differentiation, such as Rh family-C glycoprotein (RHCG), cornifelin (CNFN), kallikrein 7 (KLK7) and transglutaminase-1 (TGM1). TGM1 specifically cross-links proteins in the cornified cell envelope. The enzyme KLK7 facilitates desquamation at the epidermal surface. The differential abundance profiles are displayed in the Volcano Plot of ***Figure 5a***. Protein overlaps were considerable comparing group ‘P1-P8-P2’ to the other SCI patient group (‘P6-P9-P4-P5’): 39 of the proteins significantly increased in ‘P1-P8-P2’ showed the same directional change as in the comparison with the UTI-T cohort. That information is presented in a worksheet of Additional File 2 (Supplemental Materials).

**Fig. 5.**
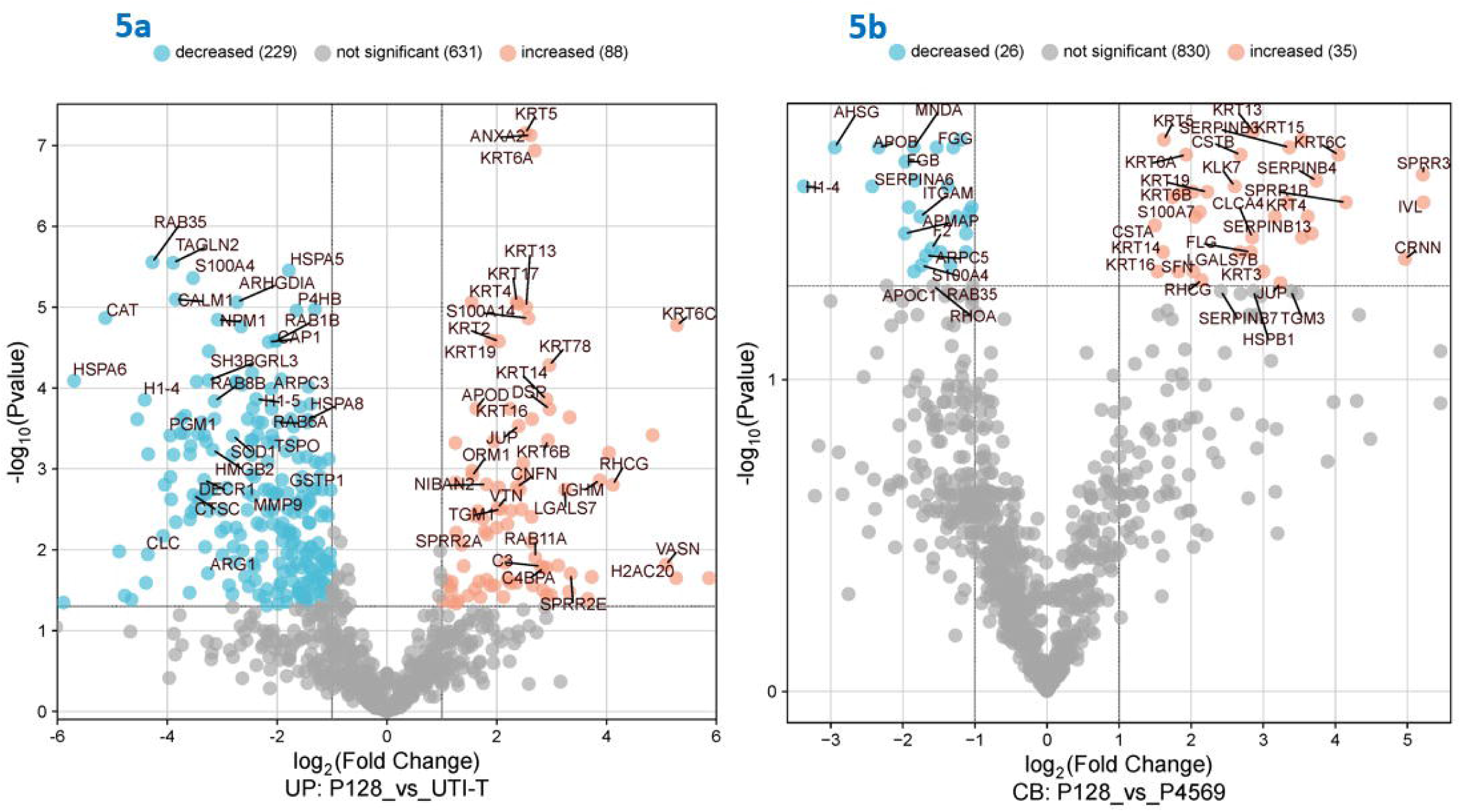
Differential abundance analyses for UP and CB proteomic datasets (a and b, respectively) visualized in Volcano plots. (a) SCI patient group ‘P2-P8-P1’ vs. UTI-T cohort; (b) SCI patient group ‘P2-P8-P1’ vs. ‘P6-P9-P4-P5’. Statistical significance thresholds were at P-value ≤ 0.02 (UP datasets) and ≤ 0.05 (CB datasets) with a fold-change set at ≥ 2. Orange-colored dots are proteins quantitatively increased in ‘P2-P8-P1’. Blue-colored dots are those decreased in ‘P2-P8-P1’ compared to the 2^nd^ group. Only a subset of dots is shown with a gene symbol.

Next, we assessed protein differences in CB proteomic datasets (group ‘P1-P8-P2’ vs. group ‘P6-P9-P4-P5’), relaxing the statistical significance in mtc•uevt-tests to a P-value ≤ 0.05. Thirty-five proteins were increased in abundance in SCI group ‘P1-P8-P2’ compared to group ‘P6-P9-P4-P5’, of which 27 were also altered in abundance in the corresponding UP dataset comparison. In support of the ‘P1-P8-P2’ group’s keratinocyte/keratinization signature, proteins increased in the CB dataset included a galectin-binding protein (LGALS7), two cystatins (CSTA and CSTB), the protein S100-A7 (synonymous to psoriasin), cornulin (CRNN) and two small proline-rich proteins (SPRR1b and SPRR3). The proteins are denoted in the plot of ***Figure 5b***. To conclude, comparative analyses are strongly supportive of the occurrence of KSM in urothelial tissues of patients 1, 2 and 8 (to a lesser extent it also pertains to P7). KRT10 was analyzed in western blots for UP and CB samples. While quantitative variability failed to deliver robust quantitative data, this cornified envelope keratin was detected as a strong 65 kDa band in western blots of several P1, P2 and P7 samples (***Figure 6***), consistent with proteomic analysis results.

**Fig. 6.**
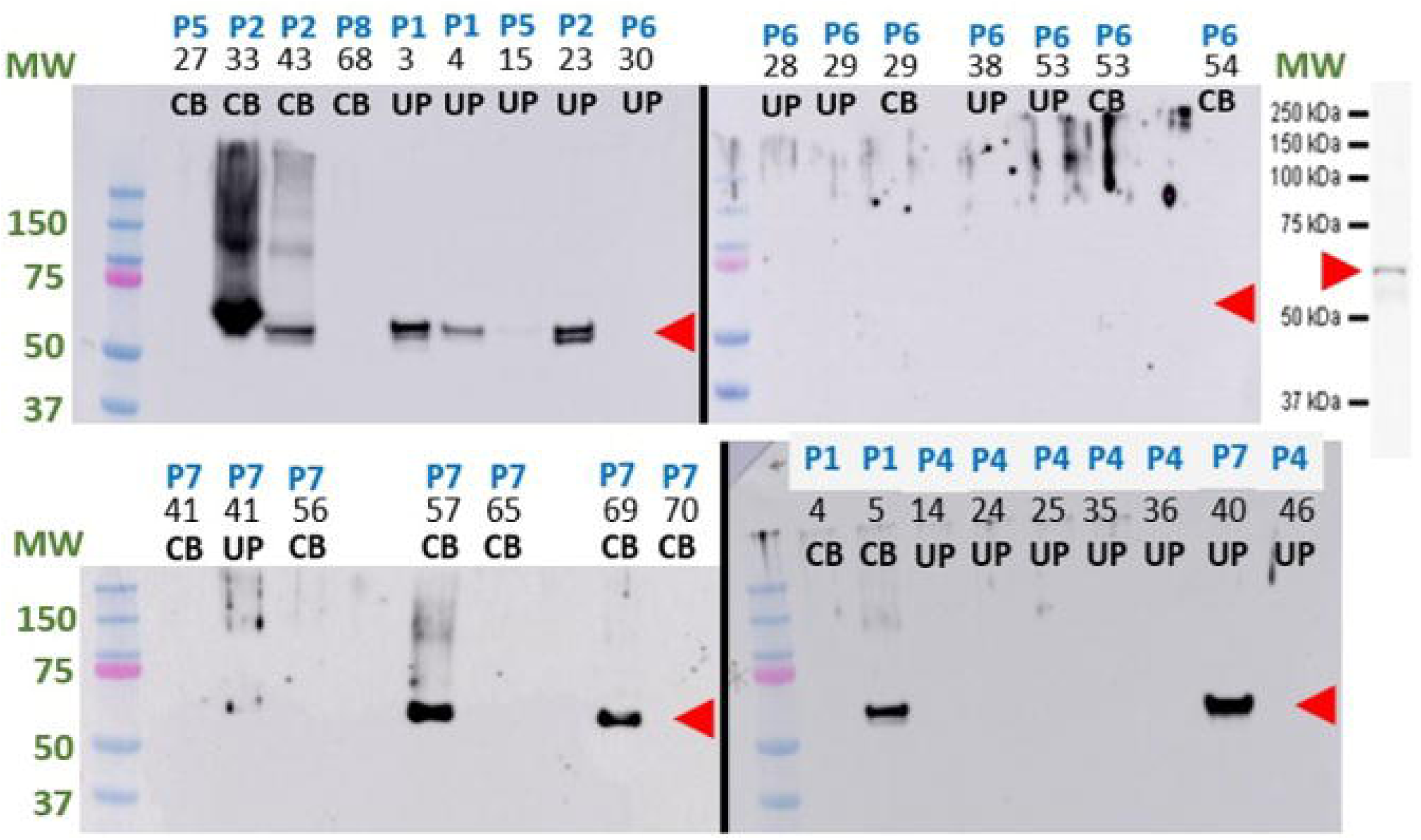
Western blots for KRT-10, highly expressed in keratinized squamous epithelial cells and the epidermal cornified envelope. KRT-10 is visualized in form of a 65 kDa band in some samples of P1, P2 and P7 but not in those of P4, P5, P6 and P8. Chemiluminescent imaging was used. Above each image, the top row denotes the patient identifier, the middle row the sample number, the bottom row the sample type. Our four blots are shown along with an image that depicts the expected band size for KRT-10 using the selected anti-KRT-10 antibody according to provider information. MW standards are denoted on the left (the pink band represents the MW value of 75 kDa). The data reveal trends for KRT-10 abundance but were not normalized between blots and not used for statistical significance assessments.

Epithelial tissue metaplasia is functionally linked to epithelial-mesenchymal transition (EMT) of cells. We found that two squamous epithelial proteins identified as EMT biomarkers, the serpins B3 and B4, were increased in abundance in SCI group ‘P1-P2-P8’ vs. ‘P6-P9-P4-P5’ (***Figure 5b***). Among the proteins decreased in abundance in UP datasets of SCI group ‘P1-P8-P2’ compared to the UTI-T group were several histones, histone-modifying enzymes and RAB GTPases. Nuclear loss is known to occur during cornified envelope formation and would explain decreased histone abundance in ‘P1-P8-P2’ datasets. Some RAB GTPases have been linked to membrane trafficking in urothelial cells and prevention of UPEC bacteria to persist in intracellular urothelial reservoirs, although the functional implications of decreased GTPase abundance in trans-differentiated bladder tissue is unclear. Some down-regulated proteins are depicted in the Volcano plots of ***Figure 5***. All 56 CB proteomic datasets with protein lists, abundance values, normalization steps and statistical analyses are presented in Additional File 5 (Supplemental Materials).

### 3.5. Protein network analysis strengthens evidence of KSM in the patient group ‘P1-P8-P2’

Proteins significantly increased in abundance in CB datasets for the SCI group ‘P1-P8-P2’ compared to group ‘P6-P9-P4-P5’ were entered into the STRING protein network analysis tool. The result was a highly connected protein network (Additional File 6, Supplemental Materials). Protein co-expression network analysis (PCNA) performed on UP proteomic datasets of the SCI patient group ‘P1-P8-P2’ versus the cohort UTI-T resulted in a network that clustered 543 proteins into 14 distinct co-expression modules with sizes ranging from 12 to 83 proteins; 587 proteins were not assignable. Module ‘eigenproteins’ were correlated with a binary trait representing SCI patient group ‘P1-P8-P2’ and cohort UTI-T. The MEturquoise co-expression module contained 83 proteins and exhibited a strong positive correlation with the SCI phenotype (r = 0.71, FDR = 1.30 x 10⁻□). Its 18 driver proteins demonstrated high module membership (kME > 0.8) and high correlation with trait SCI ‘P2-P8-P1’. The data is presented in ***Figure 7*** and Additional File 7, Supplemental Materials. The MEturquoise module is a keratin-rich network, consistent with t-test results presented in section ***3.4***. and the STRING protein network results. Bayesian inference analysis determined that keratins (most of the 18 driver proteins) were strongly associated with trait ‘P2-P8-P1’ through Bayes Factors > 10 (Additional File 7). This module featured highly significant biological process GO term enrichments shown in ***Figure 7***, among those epidermal cell differentiation (p.adjust = 2.35 x 10^−2^□) and keratinocyte differentiation (p.adjust = 1.27 x 10^−2^□).

**Fig. 7.**
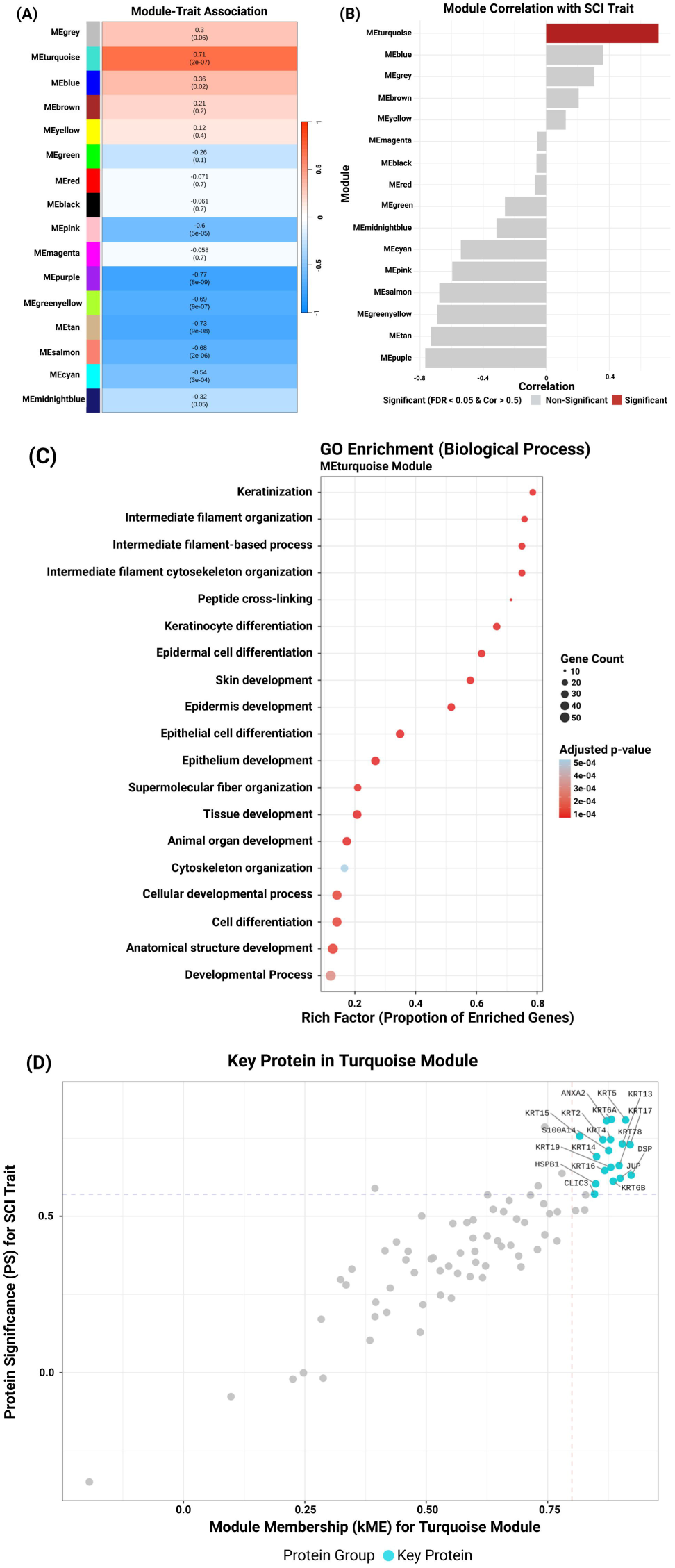
Protein Co-expression Network Analysis reveals a strong MEturquoise module featuring keratins and proteins linked to keratinocyte differentiation. With a soft-thresholding power β of 14, a network approximating scale-free topology was constructed. A. module-trait association: the algorithm clustered 543 proteins into 15 co-expression modules. B. module-SCI trait correlation: the MEturquoise module containing 83 proteins exhibited a strong positive correlation with the SCI ‘P2-P8-P1’ trait (r = 0.71, FDR = 1.30 x 10⁻). C. GO term enrichments: all highly enriched biological processes relate to keratinization and/or the cornified envelope of the epidermis. D. 18 proteins strongly associate with the trait and have a kME score > 0.8 and a significance score (PS) > 0.8.

### 3.6. Intestinal epithelial cell metaplasia in SCI patient 7 is supported by DAA and western blot data

P7 datasets were in the SCI patient group ‘P1-P8-P2-P7’ in hierarchical clustering analyses, but her UP proteomic datasets showed the greatest distance from other datasets within this group (***Figure 1***). As described in *section 3.3.*, proteins assigned to intestine-specific IC1 and IC2 clusters seemed to drive the separation of P7 samples from those of SCI patients 1, 2 and 8. DAA was used to compare P7 UP proteomic datasets with those of cohort UTI-T. Fifty-two proteins were increased in P7 datasets with a P-value ≤ 0.02 and a fold-change ≥ 2 in mtc•uevt-tests. Thirty-two of these proteins are assigned to a couple of single cell type enriched expression clusters of intestinal origin in the Protein-Atlas resource. Included are proteins secreted into the lumen of the GI tract (MUC2, MUC13, FCGBP, DEFA5, ITLN2, DMBT1, TFF2), proteins present in the cytoplasm and vesicles of enterocytes (CLCA1, ZG16), proteins localized in membranes and cell junctions of intestinal brush border cells (CDHR5, PLA2G2A, EPCAM, vilin-1) and proteins required for intestinal uptake of ions and nutrients (SLC5A1, MTTP, sucrase-isomaltase SI). A subset of these proteins is displayed in the plot of ***Figure 8a***. Twenty-five of the 32 intestinal proteins changed in abundance in comparison with the UTI-T cohort were also elevated in P7 datasets when compared to SCI group ‘P6-P9-P4-P5’. In the corresponding analysis of CB proteomic datasets (P7 vs. ‘P6-P9-P4-P5’), nine proteins were increased in P7 datasets, eight of them with cell type expression specificity for gastro-intestinal cells (the significance threshold was at P-value ≤ 0.05). This data is included in Additional File 5 (Supplemental Materials). Among these were three lectin-family regenerating islet proteins, REG1α, REG3α and REG4. Upon secretion into the gastro-intestinal lumen, REGs are active in anti-bacterial defense and tissue repair.

**Fig. 8.**
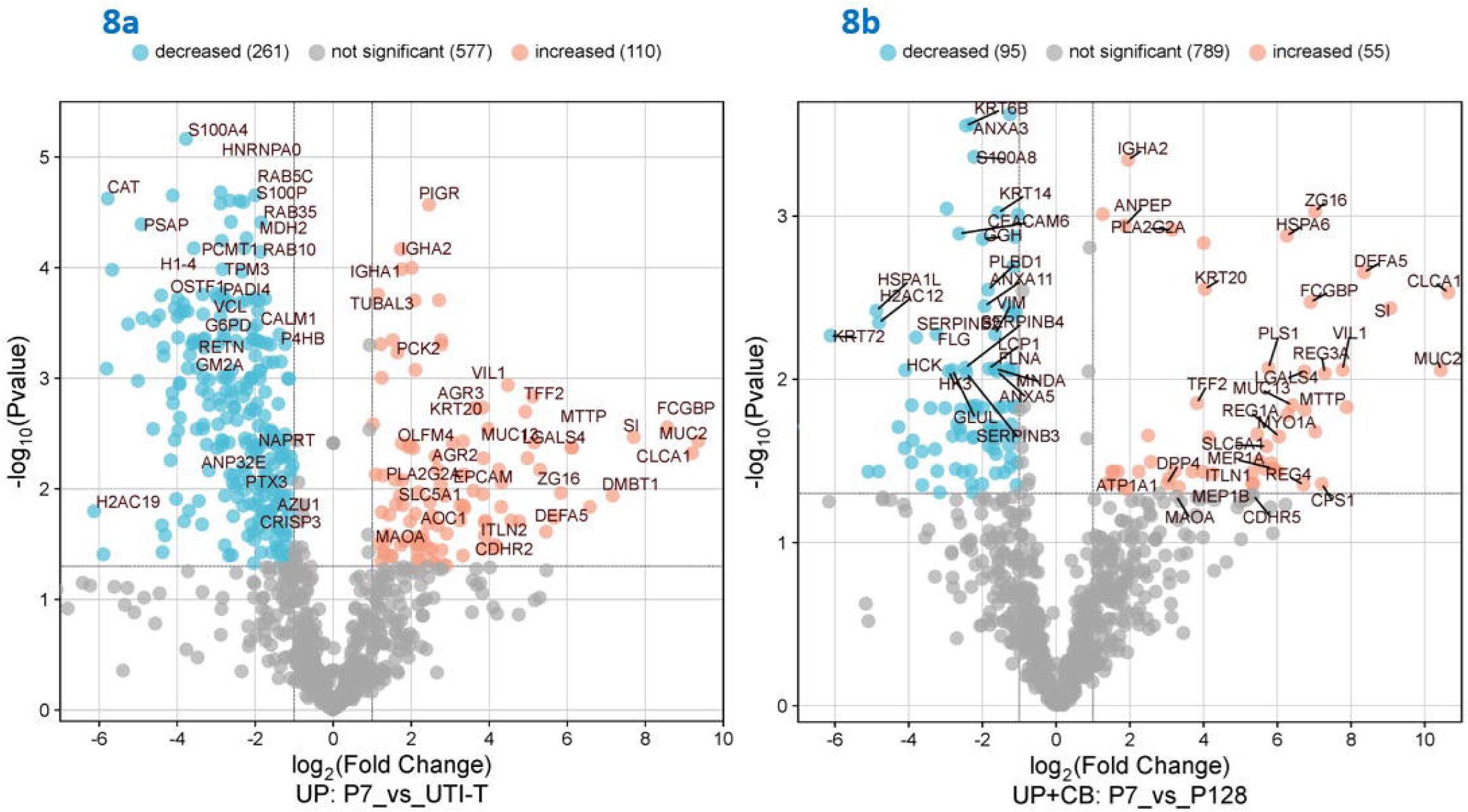
Differential abundance analyses (DAA) from proteomic datasets visualized in Volcano plots. (a.) SCI patient ‘P7’ versus UTI-T cohort (UP datasets only); (b.) SCI patient ‘P7’ versus ‘P6-P9-P4-P5’ (UP and CB datasets). Statistical significance thresholds were P-value ≤ 0.02 (for DAA in a.) or ≤ 0.05 (for DAA in b.) with a fold-change ≥ 2. Orange-colored dots are proteins increased in ‘P7’. Blue-colored dots are proteins decreased in ‘P7’ in comparison to the 2^nd^ patient group. Some proteins are denoted with their respective gene symbols.

To test the hypothesis that proteins with intestinal cell expression specificity are those that discern proteomic profiles of P7 from group ‘P1-P8-P2’, we combined UP and CB datasets in a DAA experiment. With a significance threshold at P-value ≤ 0.05 and a fold-change at ≥ 2 in mtc•uevt-tests, 56 proteins were significantly more abundant in samples from P7 (***Figure 8b***). Protein-Atlas records describe 36 of these proteins as cell type-specific to intestinal goblet cells, enterocytes and glandular epithelial cells. Using STRING protein network analysis, these proteins returned a well-connected protein network with SI, MUC-2, olfactomedin-4 and the enterocyte-specific keratin KRT-20 visualized as nodes (Additional File 8, Supplemental Materials). Cellular component GO term enrichments included the intestinal brush border and microvilli with FDRs of 1.9 10^-10^ and 3.5 10^-5^, respectively. Western blot data (***Figure 9***) show MUC-2 as a strong 120 kDa band and SI as fainter 55 and 23 kDa bands. Their detections were limited to P7 samples consistent with averaged PSM scores in proteomic datasets (1.31 and 0.42, respectively, for P7; 0 each for other SCI patient samples). In conclusion, several experiment types supported the occurrence of intestinal cell metaplasia (IM) in bladder and/or urethral tissues of SCI patient 7.

**Fig. 9.**
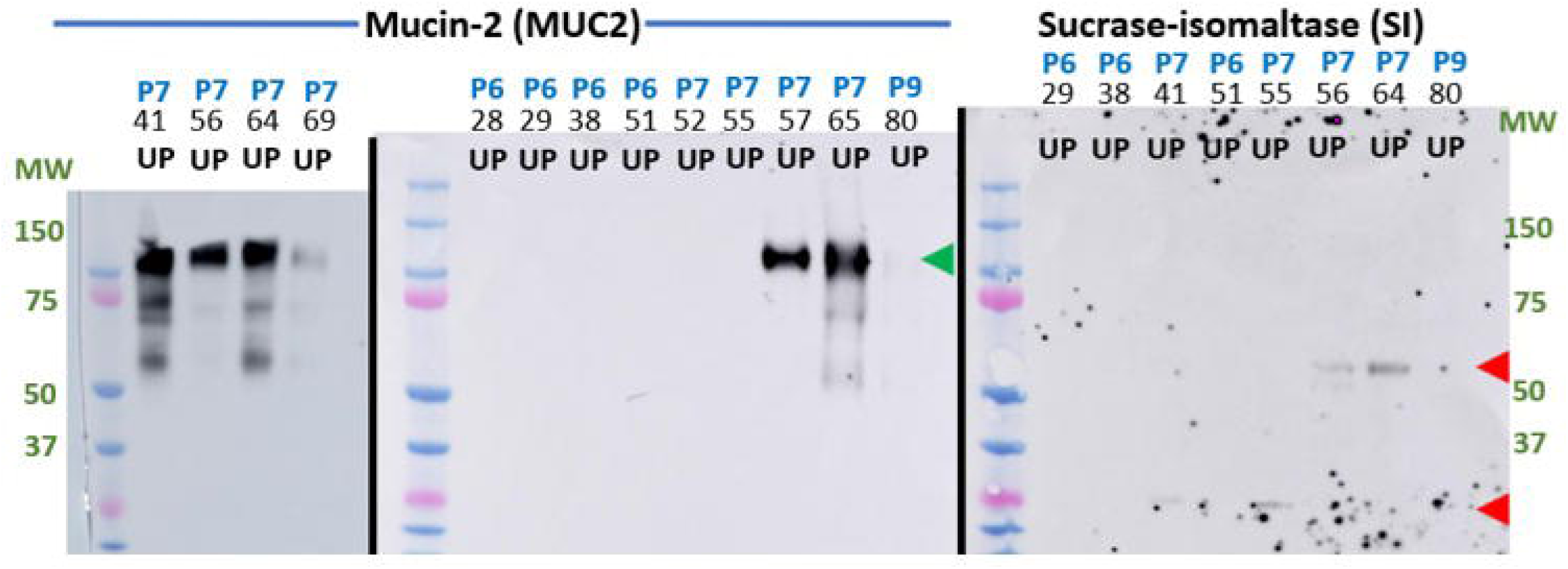
Western blots for proteins reported to be uniquely produced by intestinal epithelial cell types: mucin-2 (MUC2) and sucrase-isomaltase (SI). MUC2 is visualized as a strong 120 kDa band; SI is visualized as a set of two fainter bands (55 and 23 kDa), both only in P7 samples via chemiluminescence. Above the images, the top row denotes the patient identifier, the middle row the sample number, the bottom row the sample type (UP). Two different blots are shown for MUC2, one blot for SI. MW standards are denoted on both sides (the pink bands represent the MW values of 75 and 25 kDa). The data could not be used for statistical significance evaluations.

### 3.7. Higher complement system activation and regulation in SCI group ‘P6-P9-P4-P5’ compared to UTI-T cohort

Hierarchical clustering of proteome datasets (*section 3.3*.) featured protein clusters enriched in complement system components. Comparing SCI group ‘P6-P9-P4-P5’ with the UTI-T cohort, marked abundance increases of 16 complement factors and 19 complement regulatory proteins were noted. Proteins involved in chronic inflammation and coagulation were also more abundant in group ‘P6-P9-P4-P5’. Detailed data are included in worksheets of Additional File 2 (Supplemental Data), comparing ‘P6-P9-P4-P5’ and its subgroups (‘P4-P5’ and ‘P6-P9’) to UTI-T patient datasets. Setting a P-value threshold of ≤ 0.02 and a fold-change of ≥ 2 in mtc•uevt-tests, 75 proteins overall were differentially abundant. Entering these 75 proteins for protein network analysis (STRING), five significantly enriched biological process GO terms pertained to the complement system. FDRs were 2.8 x 10^-23^ for the classical pathway of complement activation, 5.8 x 10^-12^ for the alternative pathway and 4.1 x 10^-11^ for complement regulation. Western blot data confirmed high complement factor C3 abundance in UP samples derived from patient 4 and 5 (Additional File 9, Supplemental Materials). Integrated PCNA and Bayesian inference analysis of UP datasets (SCI vs. UTI-T patients) identified a MEred module exhibiting strong correlation with the SCI trait (r = 0.38, P-value = 6 × 10□□). Four driver proteins were complement system factors (C3 and C4B) and regulators (ITIH2 and FN1), with high module membership (kME > 0.8), high protein significance (PS > 0.8 x module-trait correlation) and strong Bayes Factor-based SCI trait association (Additional File 7, Supplemental Materials).

## Discussion

Urodynamic dysfunction and CAUTIs are life quality-diminishing pathologies that recurrently catheterized SCI patients are burdened with years or decades. The associated chronic inflammation is known to elevate long-term risk of squamous cell carcinoma (SCC) in the urinary tract. [2,33] Our longitudinal proteomic analyses of UP/CB samples from SCI patients offered the opportunity to infer histological changes as precursors of SCC or rare urothelial cancers. While not reported as frequently as in lung, cervix, esophagus and stomach tissues, both squamous cell metaplasia and IM occur in urinary tract epithelial tissues. [34] Our data suggest that KSM is relatively common in SCI patients (3 of 8 surveyed patients had a strong KSM signature). The intestinal type of *cystitis glandularis* (or IM) [18,35] was inferred from a single patient’s proteomic profiles. Only KSM is linked to elevated risk of bladder cancer risk, after a latency period of up to 28 years. [18] The KSM signature was strong in patients 1, 2 and 8 and less evident in P7 who featured the strong IM signature. Our data support the notion that non-invasive proteomic analysis of UP and CB samples is promising as a diagnostic tool to identify and differentiate subtypes of urothelial tissue metaplasia. Proteins with strong, statistically significant abundance increases in such patients are potential dysplasia risk biomarkers, e.g., KRT6A, KRT19 and S100-A14 for KSM and FCGBP, CLCA1 and MUC2 for IM.

A limitation of our work is the lack of parallel evidence from image and histological analyses obtained through cystoscopy, which could identify metaplasia or progression to dysplasia based on tissue features. From our study’s perspective, cystoscopy would have been difficult to justify due to its invasiveness, high costs including those for subsequent biopsies and the need of trained medical staff. Due to low SCC risk, cystoscopy is not routine clinical practice for frequently catheterized SCI patients. Another limitation is the fact that high throughput LC-MS proteomic analysis does not have the depth to discover oncogenic factors because those often have low expression levels and require peptide sequence IDs with specific genetic mutations. Those may be present only in rare urothelial stem and progenitor cells, further challenging detection via LC-MS when mixtures of shed bladder mucosal cells are analyzed. To gain mechanism-of-action insights on trans-differentiation of normal urothelium to tissue metaplasia and dysplasia, proteomic and genetic studies should be combined. Interestingly, though, we found an antiviral DNA cytosine deaminase, APOBEC3A, as quantitatively increased at least 5-fold in UP proteomic data of SCI group ‘P2-P8-P1’ compared to both UTI-T and ‘P6-P9-P4-P5’ datasets. This deaminase has a functional role in cancer mutations, including bladder cancers [36]. The enzyme may also contribute to long-term risk of mutations in urothelial cells of SCI patients with KSM. This discovery is an argument for further genetic and functional analyses of APOBEC3A in the context of KSM.

### Keratinizing squamous cell metaplasia

Our data supported the occurrence of KSM for two male and two female SCI patients. KSM has been reported to be more common in males, and non-keratinizing squamous cell metaplasia in females. [18] Many keratins were increased in abundance in SCI patients 1, 2 and 8 proteomic profiles (vs. UTI-T and other SCI patient groups). Some are specifically relevant to hyperproliferative keratinocytes (KRT-6a, KRT 6b, KRT6c, KRT16 and KRT-19), activated keratinocytes and SCC (KRT17), basal epidermal keratinocytes (KRT-15) and cornified envelope/advanced epidermal keratinocytes (KRT-2, KRT-1 and KRT-10). [37] Other quantitatively increased proteins were those part of cross-linked keratin filament structures (SPRRs, filaggrin, TGM1 and TGM3) or needed for keratinocyte protein breakdown and senescence (CSTA, CSTB and CLIC3). [38] Our study’s PCNA and Bayesian inference analysis supported KSM in the urinary tract of aforementioned patients by identifying specific trait-associated modules. Interestingly, DAAs from CB samples identified two serpins (B3 and B4) and kallikrein-7 as proteins increased along with keratins for the ‘P2-P8-P1’ group. Both serpins were denoted EMT biomarkers [39]. All three proteins appear to be functionally engaged in EMT. [39,40] Thus, EMT may occur prior to or along with transformation to KSM in urothelial tissues. Published data support EMT as a cellular transition necessary for development of dysplasia and early stage SCC. [41,42] These serpins have a role as peripheral blood prognostic indicators for some SSC types. [43] The important mesenchymal cell protein fibronectin 1 (FN1) was moderately increased in abundance in datasets for the group ‘P2-P8-P1’, which is consistent with transitioned mesenchymal cells in their urothelial tissues. A type-2 EMT has been associated with wound repair, a type-3 EMT with cancer progression. [44] Wound repair is clearly a key process in SCI patient urethral and bladder walls triggered by injury due to catheterization, microbial insults and immune cell infiltration. Since FN1 is critical for wound repair, we do not have support for type-3 EMT. Cystatin A (CSTA) is a protease inhibitor involved in suppressing tumor growth via EMT inhibition. We found this cystatin to be more abundant in the KSM-associated proteomic profiles. In summary, our data do not provide clarity as to whether type-3 EMT is part of the path towards metaplasia in urothelial tissues of SCI patients. The KSM we infer to have occurred in patients 1, 2 and 8 could be highly localized or broader. We observe that seven of the 25 on average most abundant proteins in UP proteomic data from those patients were keratins that contribute to cornified envelope formation. This suggests that metaplasia in the urethra and/or bladder wall is more extensive. We did not identify clinical proteomic studies related to KSM in urothelial tissues that would be useful for comparison with our data.

### Intestinal cell metaplasia

Our data yielded evidence in support of the occurrence of IM in a single female SCI patient. This metaplasia type features changes of a normal differentiated mucosal epithelium into an enteric epithelium outside of the intestinal tract and was reported to occur in the esophagus [45] and stomach. [46] An alternative term for esophageal manifestation if IM is Barrett’s esophagus, a precancerous lesion diagnosed endoscopically. [45] Normal esophageal stratified squamous epithelium is replaced by columnar epithelium with or without secretory goblet cells. [47] The etiology of gastric IM involves chronic inflammation associated with *Helicobacter pylori* infections. [48] Complete gastric IM features small intestinal-type mucosal cell trans-differentiation, containing mature absorptive cells, secretory goblet cells and a brush border. [48] This is interesting for two reasons. SCI patient 7 was also chronically infected with a bacterium (*P. mirabilis*), and its proteomic profiles featured proteins associated with each of the aforementioned enteric cell types: MUC-2 and FCGBP are of goblet cell origin; SI and MTTP are of absorptive cell origin; villin, meprin 1A and meprin 1B are of brush border cell origin. The findings tentatively support occurrence of complete IM in P7’s bladder and/or urethra. Urothelial tissues are characterized for their barrier function more than for their secretory capacity. Two secreted proteins cited in a report [49] were also identified in P7 proteomic datasets: tissue-type plasminogen activator PLAT and urokinase PLAU. Both proteins were of low abundance in P7 datasets.

While we are unaware of clinical proteomic studies linking IM to a chronically inflamed urinary tract, an impressive study examined proteomes of 155 gastric biopsies derived from patients divided in three groups: 1) mild and 2) advanced IM-like gastric lesions and 3) gastric cancers. [50]. Li *et al.* performed hierarchical clustering to discern protein signatures of these pathological groups. Of the highest interest was their comparative data analysis of biopsies linked to moderate-to-severe IM or low-grade intraepithelial neoplasia (LGIN) vs. superficial gastritis (SG). Li *et al.* identified a cluster 1 of proteins highly expressed in the IM/LGIN group that included intestine-specific proteins (MUC2, FABP1, MYO7B, ANXA13 and CDH17). In agreement with our data, FABP1 and especially MUC2 appear to be proteins of intestinal expression specificity relevant to advanced IM in urothelial and gastric mucosal cells. A cluster 2, with many gastric-specific proteins of decreased abundance in IM/LGIN vs. SG samples, contained trefoil factor TFF2. Our data revealed TFF2 increases in P7 datasets compared to those that lacked the IM signature. Gastric cancer progression biomarkers identified by Li *et al.* [50], such as APOA1BP and HPX, did not cluster with the IM signature we observed. We conclude that some features of advanced IM in gastric and urothelial mucosal tissues may be shared. Further investigations of clinical samples are needed to characterize IM occurring in different anatomical sites. Acosta *et al.* [35] surveyed 23 cases of urethral and bladder IM with a focus on oncogenic variants and histological characteristics. We did not identify proteins encoded by the mutated genes cited in that study. However, the observation of interspersed goblet cells and inflammatory cell infiltrates from histology experiments [35] was consistent with UP/CB proteomic profiles of P7.

Secretory goblet and brush border cells also occur in normal renal proximal tubular epithelia, although not as many as in the intestinal mucosa. The argument that the origin of goblet and brush border cell proteins in P7 datasets are related to exfoliation from renal tubules and leakage into urine rather than trans-differentiation of urothelial into intestinal cells. We are confident that this is very unlikely. Considerable leakage of cells from renal tubules would imply higher concentration of renal tubular proteins in urine. Cubilin, a receptor specifically expressed on renal tubular cells [51], was neither abundant in P7 datasets nor was it quantitatively increased compared to UTI-T or other SCI proteomic datasets.

### Persistent complement activation

The complement system is a double-edged sword of the immune system. While critical to detecting pathogens that cause chronic UTI and ascending renal infections, its aberrant activation is also linked to several kidney pathologies including chronic graft damage [52] and diabetic nephropathy. [53] We observed elevated complement factor levels in UP/CB proteomic datasets from SCI group ‘P6-P9-P4-P5’ compared to other patient groups. Our longitudinal profiles suggested the persistence of complement system activation but did not clarify whether its components were locally increased in expression or were more efficiently released via lymphatic or blood circulation. The possibility exists that complement activity in patients 4, 5, 6 and 9 was not increased via balancing effects of its regulators, such as inter-alpha-trypsin inhibitors and serpin G1. The latter counteract complement-mediated injury and mediate epithelial tissue repair. [54] The immune defense system’s link to metaplasia in epithelial tissues is unexplored to date.

## List of abbreviations

Abs: antibodies
CAUTI: catheter-associated urinary tract infection
CB: catheter biofilm extract
DAA: differential abundance analysis
EMT: epithelial-mesenchymal transition
FASP: Filter-Aided Sample Preparation
ID: identification
IM: intestinal cell metaplasia
KSM: keratinizing squamous cell metaplasia
mtc•uevt-tests: unequal variance multiple testing-corrected t-tests
PCA: principal component analysis
PCNA: protein co-expression network analysis
SCC: squamous cell carcinoma
SCI: spinal cord-injured
UP: urinary pellet
UTI: urinary tract infection
UTI-T: trauma patients with urinary tract infections

## Acknowledgements

This work was supported by the National Institutes of Health grant R01GM103598 titled “Urethral catheter-associated poly-bacterial biofilm formation and dispersal”. We thank Tamara Tsitrin and Yi-Han Lin for meticulous lab experiments and records tracking biological specimens and processed samples. We thank Dr. Randy Wolcott and Lisa Morrow for patient recruitment as well as collection of medical data and specimens. The funder did not take part in the study design, data collection and interpretation, or decisions to submit this work for publication.

