## Supplementary figures and images for "Keratinizing Squamous and Intestinal Metaplasia in Urothelium of Long-term Catheterized Patients with Chronic Urinary Tract Infections"

### Additional File 1

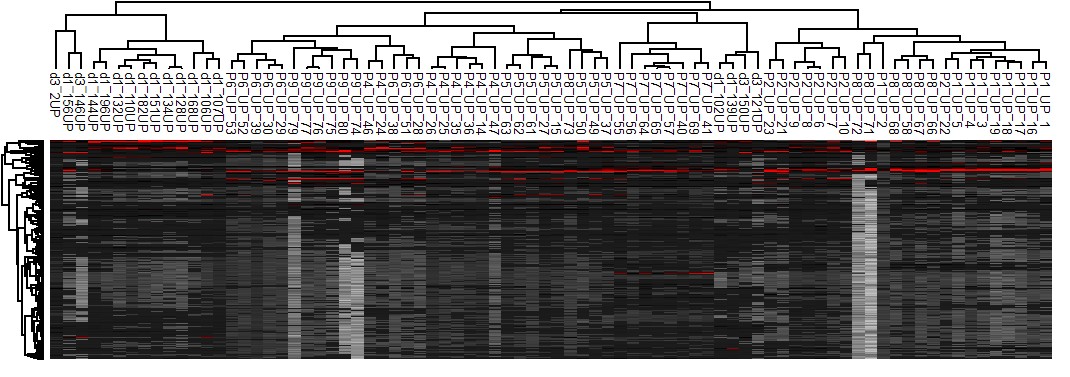

### Additional File 3

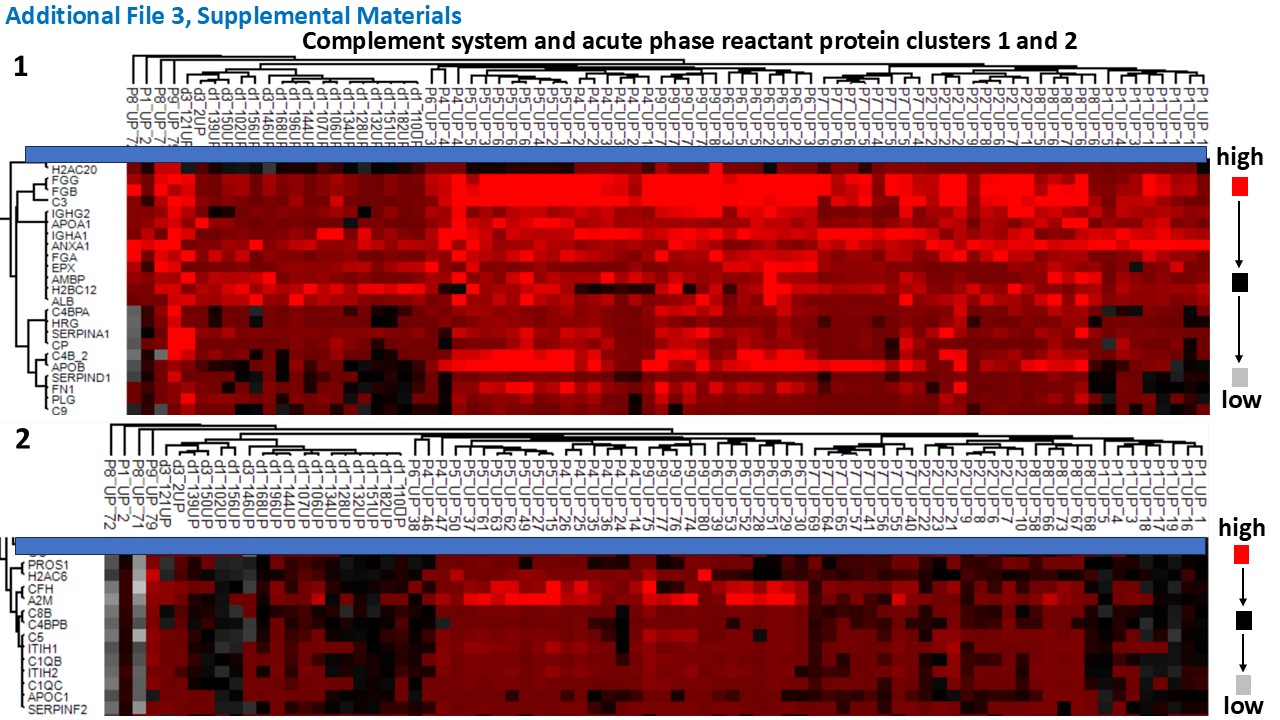

### Additional File 4

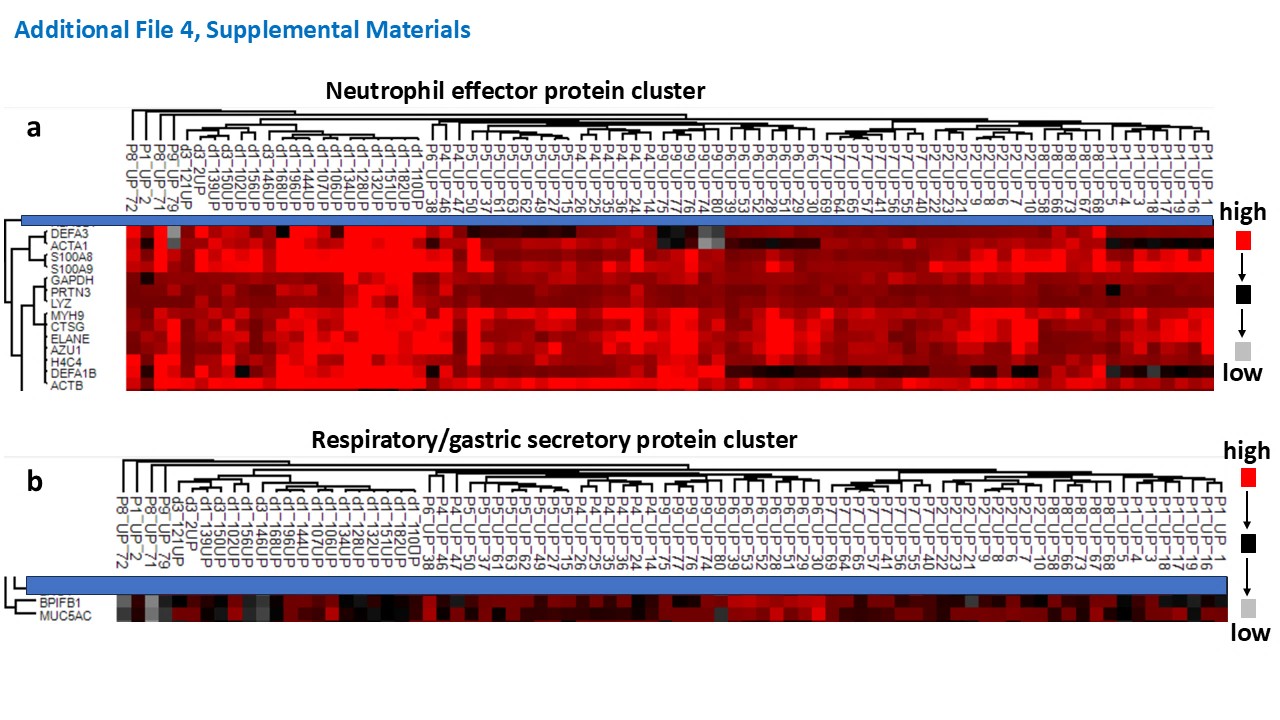

### Additional File 6

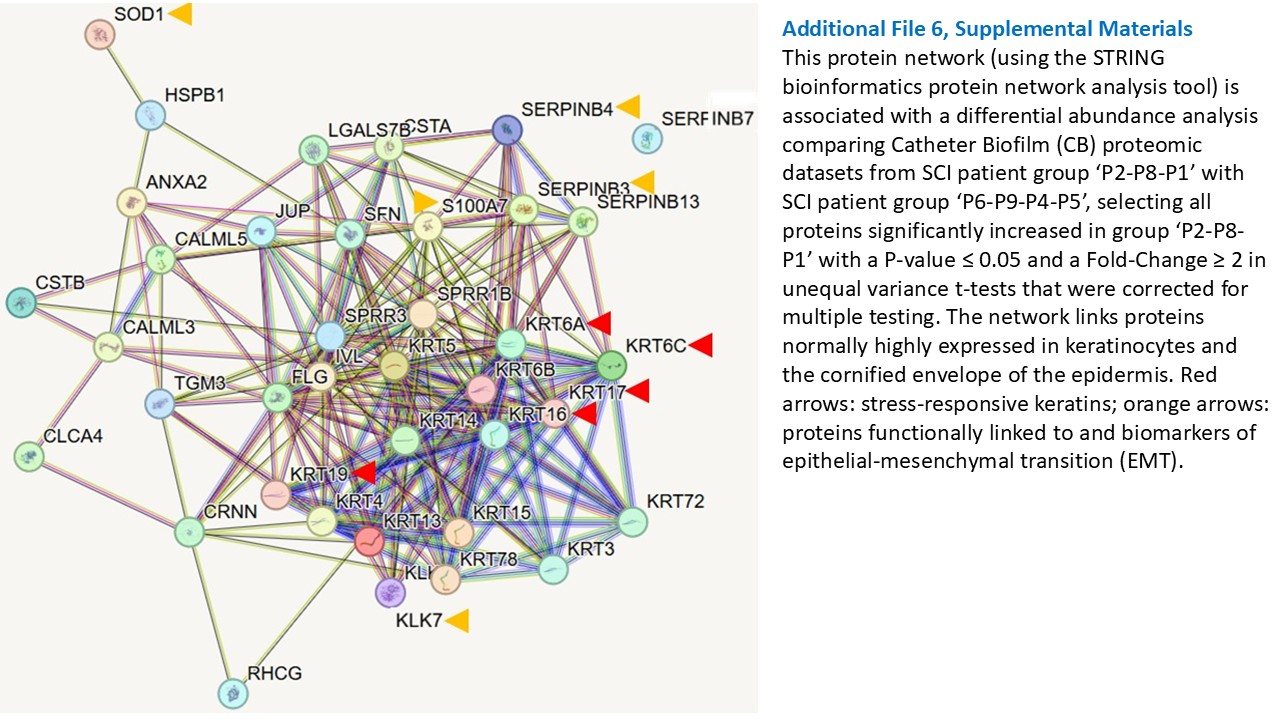

### Additional File 7

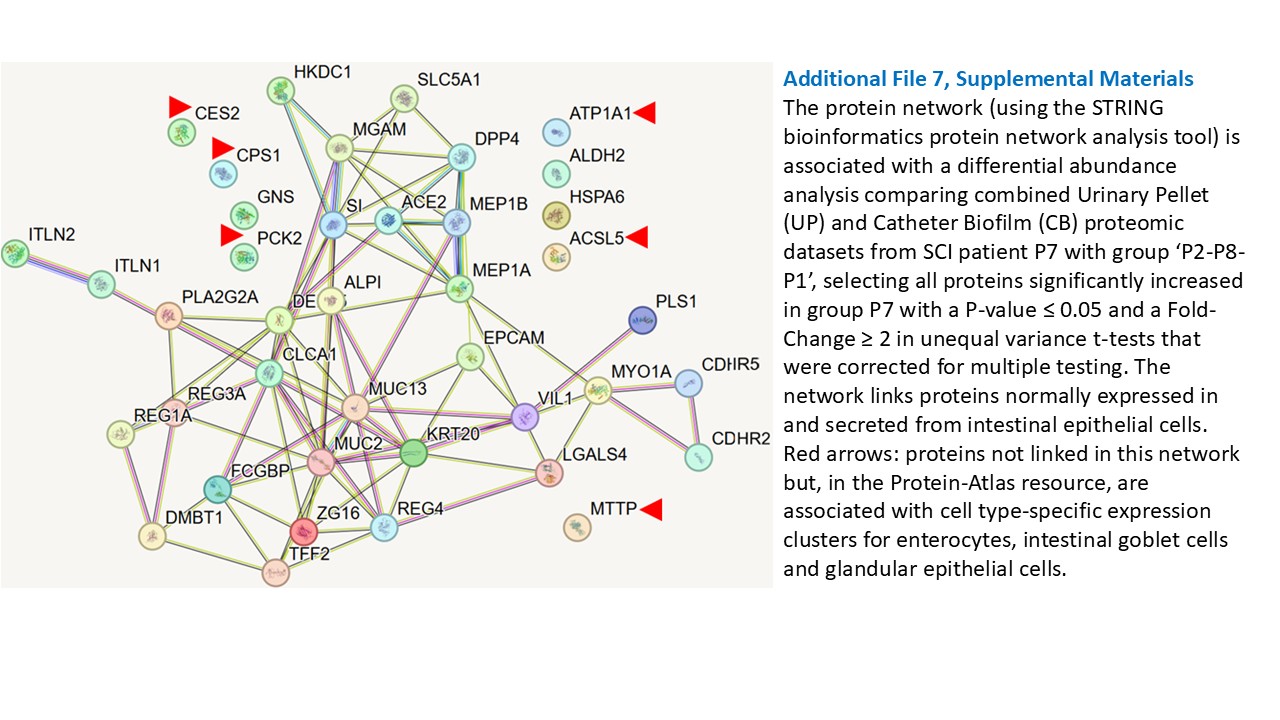

### Additional File 9

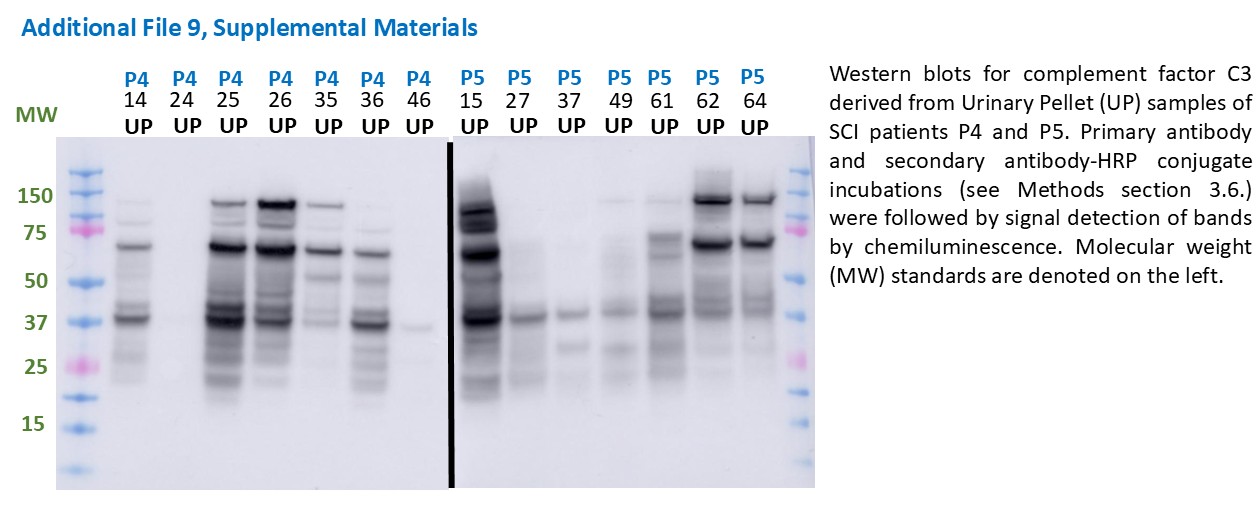
