## Additional File 10 for "Keratinizing Squamous and Intestinal Metaplasia in Urothelium of Long-term Catheterized Patients with Chronic Urinary Tract Infections"

**Supplementary Materials** (for article **“**Keratinizing Squamous and Intestinal Metaplasia in Urothelium of Long-term Catheterized Patients with Chronic Urinary Tract Infections”)

**Additional File 1.** Hierarchical clustering using Pearson correlations to cluster samples and proteins linked to UP proteomic datasets from eight longitudinally sampled spinal cord injured (SCI) patients and 18 trauma patients with acute UTIs (UTI-T cohort).

**Additional File 2.** Human proteomic datasets (137 in total from LC-MS experiments), selecting 62 Urinary Pellet (UP)-derived profiles from eight Spinal Cord Injured (SCI) patients and 19 UP-derived profiles from a cohort of trauma patients with acute Urinary Tract Infections (UTI-T).

**Additional File 3.** Two protein clusters derived from dendrogram and heat map in Figure 2 following Pearson hierarchical clustering of proteins derived from analyses of UP proteomic datasets of SCI and UTI-T patient cohorts. Complement factors and acute phase reactants were enriched in the clusters.

**Additional File 4.** Two protein clusters derived from dendrogram and heat map in Figure 2 following Pearson hierarchical clustering revealed (a) enrichments of proteins from neutrophil origin and (b) secreted effectors from respiratory/gastric epithelial cells.

**Additional File 5.** Human proteomic datasets (137 in total from LC-MS experiments), selecting 56 Catheter Biofilm extract (CB)-derived profiles from eight Spinal Cord Injured (SCI) patients.

**Additional File 6.** Protein network linked to differentially abundant proteins comparing UP proteomic profiles of SCI patient group ‘P2-P8-P1’ with those of group ‘P6-P9-P4-P5’. The network connects numerous proteins highly enriched in terminally differentiated keratinocytes and the cornified envelope of the epidermis.

**Additional File 7.** Protein co-expression network and Bayesian inference analyses comparing SCI patient groups with UTI-T cohort with description of significant modules and GO term enrichments.

**Additional File 8.** Protein network linked to differentially abundant proteins comparing UP/CB proteomic profiles of SCI patient P7 with those of group ‘P2-P8-P1’. The network connects numerous proteins unique to or at least enriched in expression in intestinal epithelial cells.

**Additional File 9.** Western blots for complement factor C3 across UP samples from SCI patients P4 and P5, with multiple C3 bands visualized via chemiluminescent staining.
